# Estimation of contemporary effective population size in plant populations: limitations of genomic datasets

**DOI:** 10.1101/2023.07.18.549323

**Authors:** Roberta Gargiulo, Véronique Decroocq, Santiago C. González-Martínez, Ivan Paz-Vinas, Jean-Marc Aury, Isabelle Lesur Kupin, Christophe Plomion, Sylvain Schmitt, Ivan Scotti, Myriam Heuertz

## Abstract

Effective population size (*N_e_*) is a pivotal evolutionary parameter with crucial implications in conservation practice and policy. Genetic methods to estimate *N_e_* have been preferred over demographic methods because they rely on genetic data rather than time-consuming ecological monitoring. Methods based on linkage disequilibrium, in particular, have become popular in conservation as they require a single sampling and provide estimates that refer to recent generations. A software programme based on the linkage disequilibrium method, GONE, looks particularly promising to estimate contemporary and recent-historical *N_e_* (up to 200 generations in the past). Genomic datasets from non-model species, especially plants, may present some constraints to the use of GONE, as linkage maps and reference genomes are seldom available, and SNP genotyping is usually based on reduced-representation methods. In this study, we use empirical datasets from four plant species to explore the limitations of plant genomic datasets when estimating *N_e_* using the algorithm implemented in GONE, in addition to exploring some typical biological limitations that may affect *N_e_* estimation using the linkage disequilibrium method, such as the occurrence of population structure. We show how accuracy and precision of *N_e_* estimates potentially change with the following factors: occurrence of missing data, limited number of SNPs/individuals sampled, and lack of information about the location of SNPs on chromosomes, with the latter producing a significant bias, previously unexplored with empirical data. We finally compare the *N_e_* estimates obtained in GONE for the last generations with the contemporary *N_e_* estimates obtained in the programmes currentNe and NeEstimator.

## Introduction

Effective population size (*N_e_*) is an evolutionary parameter introduced by Sewall Wright (Wright 1931), which determines the rate of genetic change due to genetic drift and is therefore linked with inbreeding and loss of genetic variation in populations, including adaptive potential (Franklin 1980; Jamieson and Allendorf 2012; Waples 2022). The importance of contemporary effective population size in conservation biology is increasingly recognized, and the concept implemented in conservation practice (Luikart et al. 2010; Frankham et al. 2014; Montes et al. 2016) and policy (Hoban et al. 2013; Graudal et al. 2014; Kershaw et al. 2022; O’Brien et al. 2022). For example, *N_e_* has been included as a headline genetic indicator to support Goal A and Target 4 of the Kunming-Montreal Global Biodiversity Framework of the UN’s Convention on Biological Diversity (CBD 2022), as the proportion of populations within species with *N_e_* > 500, that are expected to have sufficient genetic diversity to adapt to environmental change (Jamieson and Allendorf 2012; Hoban et al. 2020).

Contemporary *N_e_* can be estimated using demographic or genetic methods (Wright 1969; Luikart et al. 2010; Wang et al. 2016; Waples 2016; Felsenstein 2019). Demographic estimators require detailed ecological observations over time for the populations of interest (Wright 1969; Nunney 1993; Felsenstein 2019), which is not necessary for genetic estimators (Wang et al. 2016; Waples 2016). Methods that can provide *N_e_* estimates based on a single sampling point in time (Wang 2016) have become particularly popular, especially in studies focused on species for which budget and time allocated are limited, elusive species that are difficult to track and monitor (Luikart et al. 2010), and species for which information about distribution is scarce. The current biodiversity crisis and the limited resources for conservation have recently fuelled the development and application of *N_e_* estimators that rely on cost-effective, non-genetic proxy data across a wide range of species of conservation concern (Hoban et al. 2020, 2021a). Population census size, *N_C_*, has been used to infer *N_e_* when genetic *N_e_* estimates are not available, relying on the ratio *N_e_*/*N_C_* = 0.1 (where *N_C_* is the adult census size of a population) (Palstra and Fraser 2012; Frankham et al. 2014; Hoban et al. 2021b). This rule-of-thumb ratio is pragmatic for conservation (but see Fady and Bozzano 2021), as shown in application tests in different countries for different species of conservation concern (Thurfjell et al. 2022; Hoban et al. 2023). However, research needs to progress to better understand *N_e_* estimation methods and potential deviations from the ratio *N_e_*/*N_C_* = 0.1, which are expected for example across populations within species or in species with life-history traits that favour individual persistence (Jamieson and Allendorf 2012; Hoban et al. 2020, 2021b; Frankham 2021; Laikre et al. 2021; Gargiulo et al. 2023). Current genetic estimators of contemporary *N_e_* work well in small and isolated populations, which match many populations of conservation concern, but they are difficult to apply in species with a large and continuous distribution (Fady and Bozzano 2021; Santos-del-Blanco et al. 2022). In such species, genetic isolation by distance, overlapping generations, and difficulty to define representative sampling strategies can affect the accuracy of estimates of *N_C_*, *N_e_* and their ratio (Neel et al. 2013; Nunney 2016; Santos-del-Blanco et al. 2022). Plant species embody some of the features mentioned above, as they often have complex life-history traits (e.g., overlapping generations, long lifespans), reproductive systems (i.e., mixed clonal and sexual reproduction, mixed selfing and outcrossing strategies) and continuous distribution ranges (Petit and Hampe 2006; De Kort et al. 2021). Therefore, they are particularly interesting to help improve our understanding of *N_e_* estimation methods.

Genetic drift generates associations between alleles at different loci, known as linkage disequilibrium (LD), at a rate inversely proportional to *N_e_* (Hill, 1981; Waples et al. 2016). LD between loci can be used to obtain a robust estimate of contemporary *N_e_* from genetic data at a single time point, and this explains the popularity of the LD method compared to the earlier developed two-sample temporal methods (Luikart et al. 2010; Waples 2023) and the development of numerous tools for the estimation of LD*N_e_* from genetic and genomic data (Do et al. 2014; Barbato et al. 2015; Wang et al. 2016; Santiago et al. 2020). The *N_e_* estimates obtained with the LD method generally refer to a few generations back in time (Luikart et al. 2010; Do et al. 2014) and, depending on the genetic distances between loci, it is possible to obtain *N_e_* at different times in the past (Santiago et al. 2023; see also the review on timescales of *N_e_* estimates in Nadachowska-Brzyska et al. 2022). In particular, LD between closely linked loci can be used to estimate *N_e_* over the historical past (Sved 1971; Hayes et al. 2003; Qanbari et al. 2010; Do et al. 2014; Barbato et al. 2015; Wang et al. 2016; Santiago et al. 2020), whereas loosely linked or unlinked loci can be used to estimate *N_e_* in the recent past (Waples 2006a; Waples and Do 2008; Sved et al. 2013; Wang et al. 2016; Qanbari 2019). However, as other methods to estimate *N_e_*, the LD method is not devoid of biases and drawbacks, mostly relating to the assumption that the population is isolated, which is rarely satisfied (Hill 1981; England et al. 2010; Waples and England 2011; Waples 2023), and to the occurrence of age-structure in populations (Nunney 1991; Yonezawa 1997; Waples and Do 2010; Robinson and Moyer 2013; Waples et al. 2014; Hössjer et al. 2016; Ryman et al. 2019).

In this study, we aimed to explore the limitations of plant genomic datasets when estimating contemporary *N_e_*. We mostly focused on estimating *N_e_* using the software programme GONE (Santiago et al. 2020), but we also provide *N_e_* estimates obtained in NeEstimator (Do et al. 2014) and the recently developed programme, currentNe (Santiago et al. 2023). These programmes provide recent historical and contemporary *N_e_* estimates, respectively, using the LD method, though they differ mostly in the data requirement and timescales of estimates provided. GONE is the first programme using the LD method capable of exploiting the full range of LD among loci in a dataset, therefore providing *N_e_* estimates that are reliable up to 200 generations ago; NeEstimator and currentNe provide *N_e_* estimates that represent the average over few recent generations, and the exact number of generations representing an estimate increases with the number of chromosomes of the species (Santiago et al. 2023).

We explored the technical requirements of GONE by conducting power analyses aimed at testing how the number of SNPs, the proportion of missing data, the number of individuals, the lack of information about the location of SNPs on chromosomes, and the occurrence of population structure might affect *N_e_* estimation. The *N_e_* estimates obtained in GONE were then compared to the ones obtained in NeEstimator and currentNe, and discussed in light of the biological and ecological features of the species. Our findings help better understand the limitations and potentialities of genomic datasets when estimating LD-based, one-sample *N_e_*, providing new insights on how to use current methods.

## Methods

### Datasets

We selected four datasets obtained with different high-throughput sequencing techniques from different plant taxa (*Symphonia globulifera* L.f. (Clusiaceae), *Mercurialis annua* L. (Euphorbiaceae), *Fagus sylvatica* L. (Fagaceae), *Prunus armeniaca* L. (Rosaceae)), to represent different botanical groups, ecosystems, generation times and reproductive strategies. Sampling strategies in the datasets encompassed different sample sizes for markers and individuals, and datasets featured distinct levels of population genetic structure (Table 1).

**Table 1.** Details of the different plant genomic datasets analysed in the present study.

| Species name | Life-form | Reproductive system of populations analysed | Gene pools (#samples) | Data type | Average frequency of missing data per individual | #chromosomes/scaffolds/contigs analysed in GONE | Average #SNPs per scaffold or chromosome* | Total #SNPs ** | Reference | Issues explored (affecting $N_e$ estimation in GONE) |
| --- | --- | --- | --- | --- | --- | --- | --- | --- | --- | --- |
| *Symphonia globulifera* L.f. | Perennial (tree) | Monoecious, mixed mating with predominant outcrossing (Degen et al. 2004) | Species 1 (228)<br>Species 2 (107)<br>Species 3 (30) | Targeted sequence capture | 0.04 | 125 (contigs) | 247 | 30,863 | Schmitt et al. 2021 | Minimum number of SNPs required |
| *Mercurialis annua* L. | Annual | Various mating systems, analyses based on dioecious populations; obligate outcrosser (González-Martínez et al. 2017) | Atlantic (12)<br>Core (16)<br>Mediterranean (12) | Targeted gene (exome) capture | 0.01 | 48 (contigs) | 670 | 32,151 | González-Martínez et al. 2017 | Influence of sample size |
| *Fagus sylvatica* L. | Perennial (tree) | Monoecious, predominant outcrossing (Merzeau et al. 1994) | Mt. Ventoux, France (167) | Whole genome sequencing | 0.81 (with 27 scaffolds) | 12-150 (scaffolds) | ~470K (with 27 scaffolds) | ~13 M (with 27 scaffolds) | See data availability section | Influence of missing data |
| *Prunus armeniaca* L. | Perennial (tree) | Monoecious, self-incompatible (Groppi et al. 2021) | Southern (56)<br>Northern (199) (see Supplementary Table 1) | Whole genome sequencing | 0.07 | 8 (chromosomes) | ~3 M (440K) | ~24 M (3.5 M in the subsampled dataset) | Groppi et al. 2021 | Influence of number of SNPs, of missing data, of sample size, of population structure, of using scaffolds instead of chromosomes |

\*in the map file, number of lines divided by number of scaffolds/chromosomes;
\*\*number of lines in the map file

For boarwood, *S. globulifera* s.l., a widespread and predominantly outcrossing evergreen tree typical of mature rainforests in Africa and the Neotropics (Degen et al. 2004; Torroba-Balmori et al. 2017), we used the targeted sequence capture dataset described in Schmitt et al. (2021). Three sympatric gene pools were identified in a lowland forest in French Guiana, likely corresponding to three biological species, described as *Symphonia* sp. 1, *Symphonia* sp. 2 and *Symphonia* sp. 3 (Schmitt et al. 2021). To avoid the influence of admixture on the estimation of *N_e_*, we first divided the dataset in three subsets based on the analysis of genetic structure performed in the software Admixture v1.3.0 (see Schmitt et al. 2021), selecting only the individuals with a Q-value (cluster membership coefficient) ≥ 95% to each of the three genetic clusters (Species 1, Species 2 and Species 3; Supplementary File 1). We then selected the 125 genomic scaffolds with the largest number of SNPs (see Table 1).

For the annual mercury, *M. annua*, an annual plant with variable mating systems (monoecious, dioecious, androdioecious), ploidy levels (2x, 4x-12x) (Obbard et al. 2006b, a), potential to produce seed banks, and typical of open or disturbed habitats in Europe and North Africa, we used the gene capture data set described in (González-Martínez et al. 2017), obtained from 40 diploid dioecious individuals grown from seeds, representative of ten localities and three main gene pools in the species (as described after the fastStructure analysis in González-Martínez et al. 2017). We selected the 48 scaffolds with the largest number of SNPs and ran the analyses by considering separately each gene pool: (1) ancestral populations from Turkey and Greece (“Core”), (2) range-front populations from northeastern Spain (“Mediterranean”), or (3) range-front populations from northern France and the UK (“Atlantic”) (see Table 1).

For the common beech, *F. sylvatica*, a deciduous predominantly outcrossing tree of European temperate forests (Merzeau et al. 1994), we analysed genomic scaffolds from a single, contiguous stand (plot N1; (Oddou-Muratorio et al. 2021)) within a relatively isolated French population (Mt. Ventoux, southeastern France), in which population genetic structure is neither observed nor expected (Csilléry et al. 2014). Mapping of short-reads paired Illumina sequences was independently performed for each one of the 167 individuals of the population against the genome assembly (available at www.genoscope.cns.fr/plants) using bwa-mem2 2.0 (Li and Durbin 2009). SNPs were first called using GATK 3.8 (Van der Auwera and O’Connor 2020) using the following parameters: -nct 20 -variant_index_type LINEAR variant_index_parameter 128000. SNPs were also called using samtools v1.10 / bcftools v1.9 (Danecek et al. 2021) with default parameters. Following these two SNPs calling steps, we performed a three-steps filtering process: (i) only diallelic SNPs were kept, (ii) the minimum allele frequency (MAF, upper case used at the individual level), calculated on the basis of all the reads containing the SNP, was set to 30% (note that GONE does not require the application of MAF filtering, and such filtering might cause a small upward bias in the estimation), (iii) individual genotypes with sequencing depth less than 10 were recoded into « ./. » meaning that both alleles are missing. We then identified SNPs found by both GATK and samtools using the - diff flag of vcftools v0.1.15 with tabix-0.2.5 (Danecek et al. 2011). A nucleotide polymorphism was considered to be a SNP if at least one individual was found to be heterozygous at the position. On average, for each individual, 88.5% of the sequencing reads mapped properly onto the assembly. The final VCF contained 18,192,174 variants, and is available at the Portail Data INRAe (doi:10.57745/FJRYI1).

We re-ordered the 406 genomic scaffolds available based on their number of SNPs, and selected 150 scaffolds with the largest number of SNPs. We tested different combinations of input subsets, with numbers of scaffolds ranging from 12 to 150 (provided that SNPs per scaffold < 1 million and total number of SNPs < 10 millions, see the requirements of GONE below), and numbers of individuals ranging from 5 to 167 (total sample size).

For the apricot, *P. armeniaca*, we estimated *N_e_* using whole genome resequencing data (21× depth of coverage by ILLUMINA technology) for wild Central Asian, self-incompatible populations of the species (Groppi et al. 2021). Variant sites were mapped to the eight chromosomes of the species and ranged between 2.3 and 6.2 million per chromosome (total number of variant sites: 24 M). As these exceeded the total number allowed in GONE, we downsampled the number of SNPs prior to the analyses. We also analysed the datasets by considering the different gene pools recovered in Groppi et al. (2021) (Supp. Fig. S20), namely the Southern (red cluster) and Northern (yellow cluster) gene pools, as obtained in fastStructure (Raj et al. 2014) (see next subsection).

### Data analyses in GONE

#### Analyses for all species

We performed *N_e_* estimation in the software GONE (Santiago et al. 2020). GONE generates contemporary or recent historical estimates of *N*e (i.e., in the 100-200 most recent generations) using the LD method. GONE requires linkage information, ideally represented by SNPs mapped to chromosomes. Chromosome mapping is rarely available for non-model species, and in our case was only fully available for the apricot (*P. armeniaca*) dataset. In the absence of chromosome mapping information for the other species, we treated genomic scaffolds as chromosomes. In terms of requirements, GONE accepts a maximum number of chromosomes of 200 and a maximum number of SNPs of 10 million, with a maximum number of SNPs per chromosome of 1 million, although the software uses up to 50,000 random SNPs per chromosome for the computations when the total number of SNP is larger. A complete workflow of the analyses carried out in GONE is available at https://github.com/Ralpina/Ne-plant-genomic-datasets (Gargiulo, 2023); the input parameter file used for the final analyses is available in Supplementary File 2.

#### Influence of missing data on N _e_ estimation

The influence of missing data on *N_e_* estimation in GONE was evaluated using the dataset from *F. sylvatica* . After keeping 67 individuals with less than 95% missing data, we permuted individuals (without replacement) to generate 150 datasets of 35 individuals, and estimated *N_e_* in GONE for each dataset. Proportion of missing data per individual for each permuted dataset was calculated in vcftools v0.1.16 (Danecek et al. 2011) from an average of ∼25% to 95%; results were plotted in R v4.2.2 (R Core Team 2019). In addition, we used the dataset of *P. armeniaca* to evaluate how *N_e_* changed when manually introducing missing data. We selected all individuals from the Northern gene pool with a Q-value (cluster membership coefficient) ≥ 99% (77 individuals) to rule out the influence of admixture, and replaced some of the individual genotypes with missing values using a custom script (available at: https://github.com/Ralpina/Ne-plant-genomic-datasets). We generated two datasets with a proportion of missing data per individual of 20% and 40%, respectively, and then computed *N_e_* in GONE for each dataset obtained.

#### Influence of number of SNPs on N_e_ estimation

The influence of the number of SNPs on *N_e_* estimation in GONE was evaluated using the dataset of *P. armeniaca* . From the Northern gene pool, we first selected the individuals with a Q-value ≥ 99% to rule out the influence of admixture. We drew random subsets of variant sites (without replacement) including 40K, 80K, 150K, 300K, 500K, 3.5M, 7M, and 10M SNPs, respectively, and generated 50 replicates for each subset; we then estimated *N_e_* in GONE for each subset and obtained the geometric mean and the 95% confidence intervals across the 50 replicate subsets with the same number of SNPs (using the functions *exp(mean(log(x)))* and *quantile* in R).

#### Influence of sample size on N_e_ estimation

We used the Northern gene pool of *P. armeniaca* to assess how *N_e_* estimates changed depending on the number of samples considered and the uncertainty associated with individual sampling. We first downsampled the number of SNPs to 3.5M (to satisfy GONE requirements), and varied the sample sizes included in the analyses from 15 to 75 (i.e., approx. the total number of individuals of the Northern gene pool with a Q-value ≥ 99%). For each sample size group, we generated 50 subsets (without replacement within the subset) of individuals and estimated *N_e_* in GONE for each subset; we then estimated the geometric mean and the 95% confidence intervals across subsets with the same sample size (using the functions *stat_summary(fun.data = median_hilow, fun.args = list(conf.int = 0.95)* and *stat_summary(fun = “geometric.mean”* (psych package) in R).

#### Influence of population admixture on N _e_ estimation

We also evaluated how genetic structure within gene pools influenced *N_e_* estimation in GONE for both the Southern and Northern gene pools of *P. armeniaca*. We first downsampled the number of SNPs to 3.5M to satisfy GONE requirements, as described above. We then distributed the individuals of each gene pool into five (overlapping) subsets based on individual Q-values (lower bounds of 70%, 80%, 90%, 95%, and 99%), resampled individuals (without replacement) in each Q-value subset 50 times, standardising sample sizes to the sample size of the smallest Q-value subset within a gene pool (i.e., 21 individuals as in the 99% Q-value subset of the Southern gene pool and 77 individuals as in the 99% Q-value subset of the Northern gene pool, see Supplementary Table S1 for original sample sizes). We then estimated *N_e_* in GONE and obtained 95% confidence intervals across the 50 resampled datasets of the same Q-value subset within a gene pool (using the R function *stat_summary* mentioned above). We also combined all individuals from the two gene pools (255 individuals), resampled 77 individuals 50 times without replacement, and estimated *N_e_* in GONE and the related confidence intervals as explained above.

#### Effect of using genomic scaffolds rather than chromosomes

We evaluated the effect of using genomic scaffolds to estimate linkage groups when chromosome information is not available. Using the downsampled dataset of 3.5M SNPs from *P. armeniaca*, we selected from the Northern gene pool 45 random individuals with a Q-value ≥ 99%, to rule out the influence of admixture. For this dataset, five different chromosome maps were then created, progressively assigning SNPs to 8 (true value), 16, 32, 64 and 128 chromosomes (as if they were genomic scaffolds, see script and related explanation at https://github.com/Ralpina/Ne-plant-genomic-datasets#4-effect-of-using-genomic-scaffolds-instead-of-chromosomes-on-ne-estimation). We then estimated *N_e_* in GONE using five corresponding chromosome map files and keeping the same ped (genotypes) file.

### Data analyses in NeEstimator

We also used the LD method as implemented in the software NeEstimator v2 (Do et al. 2014) to estimate *N_e_* in our datasets. NeEstimator assumes that SNPs are independently segregating (typically, SNPs at short physical distances, for example those in the same short genomic scaffolds or loci, are filtered previous to analysis, see below), and therefore it provides an *N_e_* estimate based on the LD generated by random genetic drift, which reflects *N_e_* in very recent generations (Waples et al. 2016). However, accuracy and precision will be both affected by (1) the assumption of independent segregation in genomic data sets, as SNPs are necessarily packed on a limited number of chromosomes and thus they provide non-independent information, and especially (2) the occurrence of overlapping pairs of loci, each locus appearing in multiple pairwise comparisons (i.e., two aspects of the issue known as pseudoreplication; (Purcell et al. 2007; Waples et al. 2016; 2022; Waples 2023)). Although the influence of this issue on bias and precision is difficult to address completely, some bias corrections have been proposed, for example applying a correction based on the genome size of the species being analysed (formula in Waples et al. 2016), restrict comparisons to pairs of loci occurring on different chromosomes (Waples 2023), or using only one SNP per scaffold or thinning scaffolds based on discrete window sizes (Purcell et al. 2007). To adjust for the bias, we therefore applied the correction in Waples et al. (2016), by dividing the *N_e_* estimates obtained by *y=0.098+0.219 × ln(Chr)*, where *Chr* is the haploid number of chromosomes, when information about the number of chromosomes was available.

As low-frequency alleles upwardly bias *N_e_*, we followed the recommendations in Waples (2023) and excluded singleton alleles (Waples and Do 2010; Waples 2023). We also ran the analyses without applying a filter for rare alleles, to be able to compare the results obtained in NeEstimator with those from GONE and currentNe. Confidence intervals were obtained via jackknifing over samples (Do et al. 2014; Jones et al. 2016). As NeEstimator cannot handle very large datasets (with > 100,000 loci, see https://www.molecularfisherieslaboratory.com.au/neestimator-software/), we reduced the number of SNPs in the *F. sylvatica* and *P. armeniaca* datasets by randomly subsampling 50,000 SNPs across chromosomes.

### Data analyses in currentNe

We used the newly developed software programme currentNe (Santiago et al. 2023) to obtain contemporary *N_e_* estimates that are directly comparable to the ones obtained in NeEstimator (referring to the most recent generations in the past). The practical advantages of currentNe are the possibility to include thousands of SNPs in the analyses (with an upper limit of 2 million loci), the lack of a minor allele frequencies requirement, and the lower computational effort. Moreover, the software produces confidence intervals around *N_e_* based on artificial neural networks, can accommodate complex mating systems and is accurate with small sample sizes (Santiago et al. 2023). We estimated *N_e_* in currentNe for all the species included in our study except *S. globulifera* s.l., as the software requires the number of chromosomes or the genome size in centiMorgans, which were not available for the species.

## Results and Discussion

### Data analyses in GONE

Our study explores the limitations associated with genomic datasets when estimating *N_e_* using the LD method as implemented in the programme GONE, and compares estimates of recent historical *N_e_* obtained in GONE with estimates of contemporary *N_e_* as obtained in NeEstimator and currentNe. Below, we will first focus on the limitations of plant genomic datasets as explored using the software GONE and then discuss the differences observed when *N_e_* was calculated using GONE, NeEstimator and currentNe.

One limitation usually associated with reduced representation sequencing datasets is the short length of the reads or scaffolds. We tested how this limitation would influence *N_e_* estimation in GONE using the datasets of *S. globulifera* and *M. annua*. The estimation of *N_e_* in GONE failed for the three biological species of *S. globulifera*, as the software returned the error “too few SNPs” for each of the three species datasets. This was caused by the relatively small number of SNPs per scaffold (averaging ∼250 SNPs) and, in turn, by the relatively short length of the scaffolds (length ranging from 5,421 to 931 positions) which prevented GONE from producing reliable *N_e_* estimates. *N_e_* estimates were instead obtained for *M. annua*, whose average number of SNPs per contig was 670 (Table 1).

#### Influence of missing data on N_e_ estimation

The effect of missing data on *N_e_* estimation is evident from the results obtained when analysing the dataset of *F. sylvatica*, and from the results obtained when analysing the dataset of *P. armeniaca* in which genotype data were manually excluded. For *F. sylvatica*, 35 individuals had a proportion of missing data < 50% (Fig. 1B). Increasing the proportion of missing data in the permuted datasets of 35 individuals produced acute increases in *N_e_* estimates in GONE (see Fig. 1A); for instance, increasing the median proportion of missing data per individual from 25% to 35% produced *N_e_* estimates increasing from 200 to 3 millions. Likewise, when missing data proportion per individual of *P. armeniaca* increased above 20%, we obtained *N_e_* estimates that were > 350 times larger than those obtained from the original dataset (average missing data proportion per individual ∼ 8%) (Fig. 2). This relationship between missing data and *N_e_* estimates is consistent with what was previously found (e.g., Marandel et al. 2020), although the loss of accuracy in the *N_e_* estimation is extreme and suggests that either individuals with > 20% missing data should be removed from the dataset before estimating *N_e_* or SNPs with missing data in a given percentage of individuals (e.g., 50% by default assumed by GONE) should be removed, provided that the dataset includes a sufficient number of SNPs. However, in species with large effective population sizes, reducing the sample size (S) to a number << true *N_e_* introduces further uncertainties in the *N_e_* estimation using the LD method, regardless of the number of loci used (Marandel et al. 2019; Waples 2023), in addition to the sampling error already expected because of the finite sample size (e.g., Peel et al. 2013).

**Figure 1.**
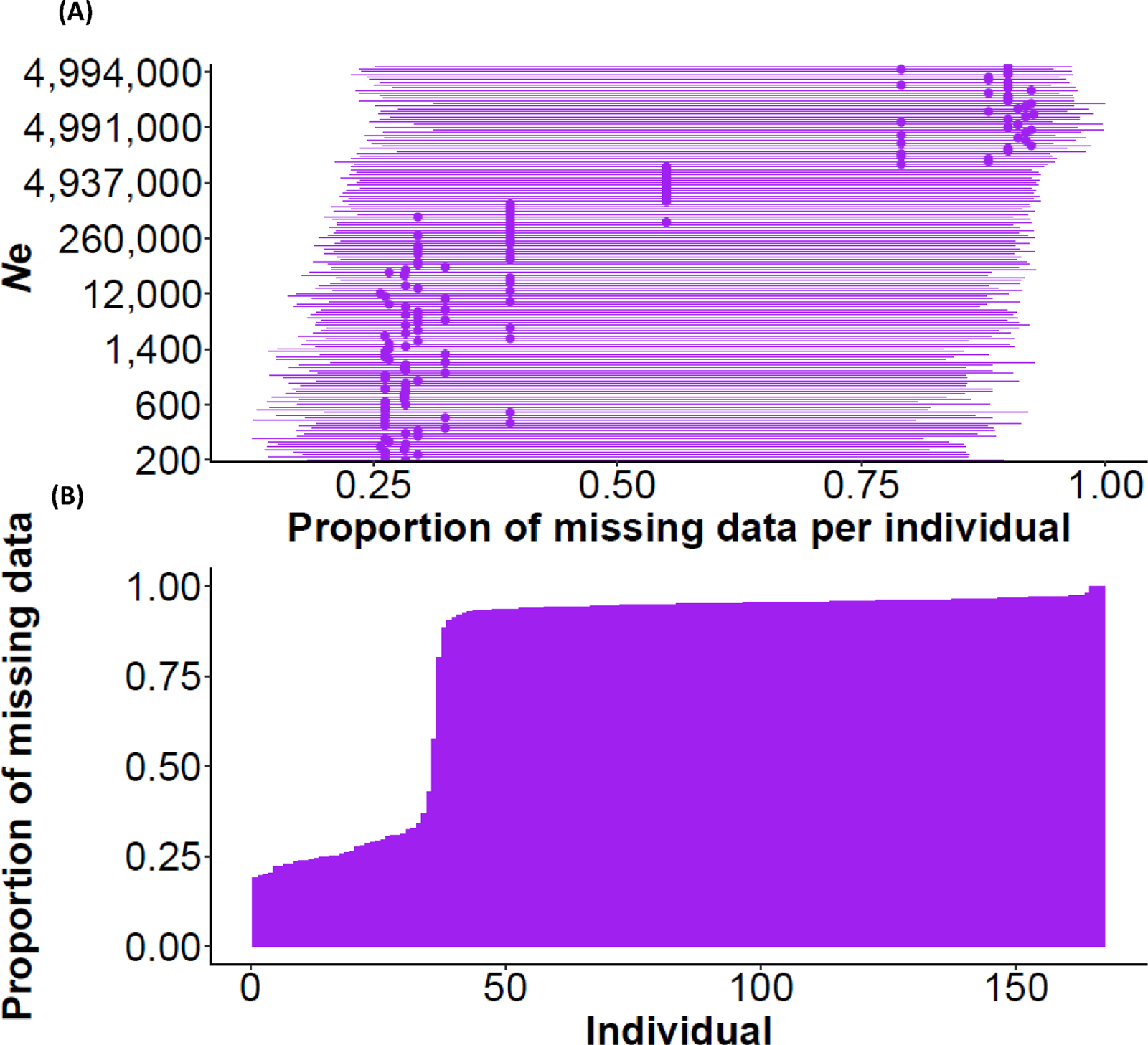
In (A), ranked median *N*_e_ estimates in the most recent generation in 150 datasets of 35 individuals with different proportions of missing data (excluding individuals with a proportion of missing data > 0.95) of *F. sylvatica*; ranges represent standard deviations for the proportion of missing data per individual. Analyses based on the dataset with the twenty-seven genomic scaffolds with the largest number of SNPs (excluding the scaffolds with > 1 M SNPs). In (B), proportion of missing data per individual in the complete dataset of *F. sylvatica*.

**Figure 2.**
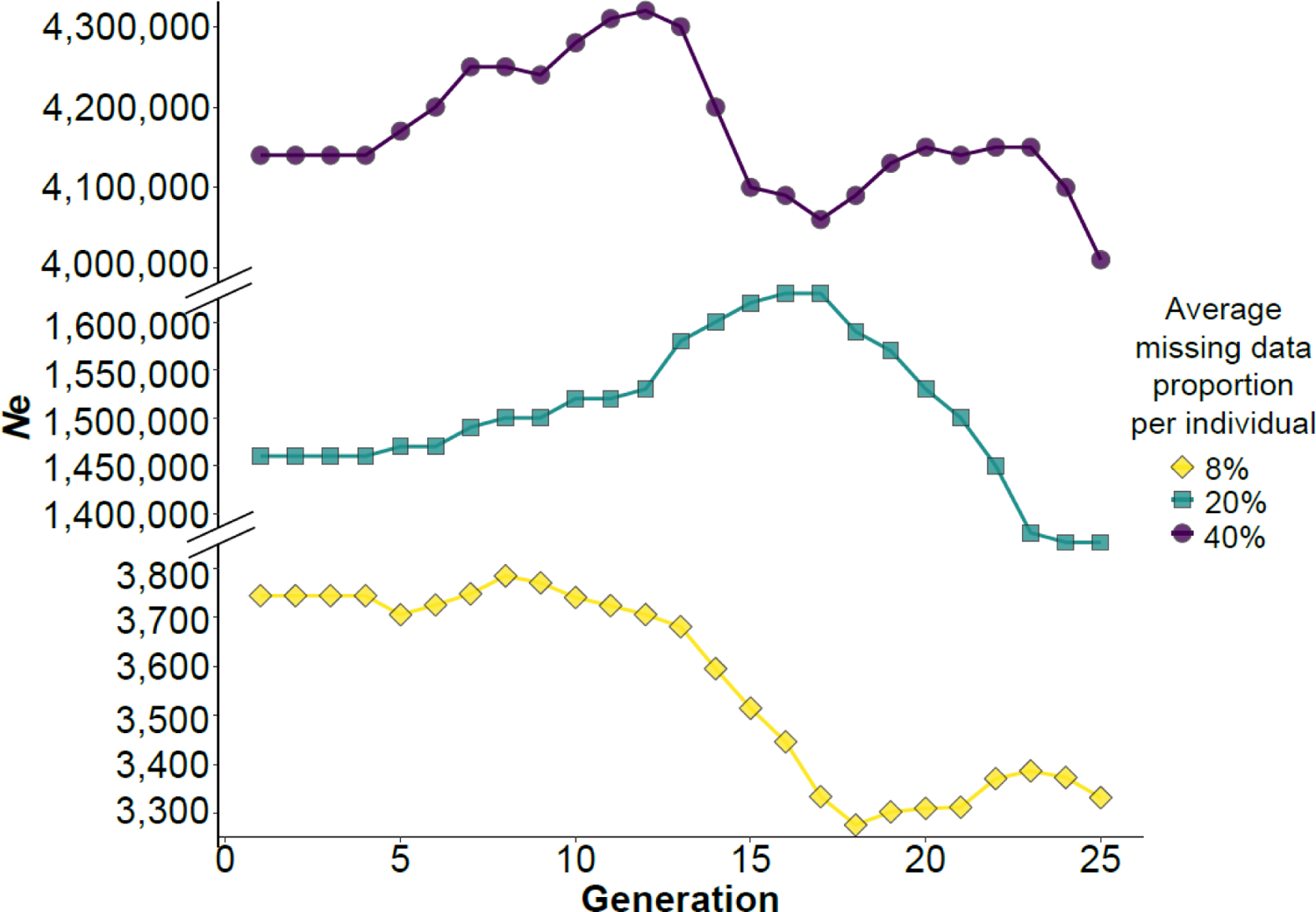
Influence of missing data on *N*_e_ estimation in GONE. Missing genotypes were manually introduced in the dataset of *P. armeniaca*, generating pseudo-genotypes with an average proportion of missing data ranging from 20% to 40%. The original dataset is shown for comparison (missing data = 8%). Note the different y-scales in the three facets.

#### Influence of number of SNPs on N_e_ estimation

The influence of the number of SNPs per chromosome was explored using the dataset from P. armeniaca (Northern gene pool), which was the only dataset with SNPs fully mapped to chromosomes. Increasing the number of SNPs per chromosome affected point N_e_ estimates only slightly, and influenced the apparent precision of the estimates more obviously, especially for a total number of SNPs above 300,000, corresponding to an average of 10,000 SNPs per chromosome of P. armeniaca used by GONE (Fig. 3). Accuracy and precision of N_e_ estimates based on LD are expected to be affected by two types of pseudoreplication: (1) the non-independent information content provided by thousands of linked SNPs, and especially (2) the occurrence of overlapping pairs of loci, each locus appearing multiple times in pairwise comparisons (Waples et al. 2016; 2022). Therefore, the narrower confidence intervals we obtained when increasing the number of SNPs are partially due to the inclusion of overlapping pairs of loci for the N_e_ estimation, which artificially increases the degrees of freedom that make CIs tight. The drop in the N_e_ geometric mean value associated with the dataset with >20,000 SNPs might be due to the inclusion of more physically linked SNPs, but it might also be due to the uncertainty associated with the specific SNPs included in the analysis.

For practical purposes, our results show that adding more than 2,000 polymorphic SNPs per chromosome, with a large sample size (∼75), does not substantially improve the accuracy and the precision of the estimation, in line with what is shown in previous studies focusing on LDN_e_ (Marandel et al. 2020). Santiago et al. (2020) noted that the accuracy of the estimation is proportional to sample size and to the square root of SNPs pairs, and therefore researchers might partially compensate for small sample sizes by increasing the number of SNPs. However, as the information content of a dataset depends on the amount of recombination and on the pedigree of the individuals included in the analyses, an estimation based on a small number of samples will not necessarily be representative of the entire population, especially if N_e_ is large (King et al. 2018; Santiago et al. 2020; Waples 2023). Furthermore, the marginal benefit of increasing the number of SNPs beyond tens of thousands is counterbalanced by poor precision if CIs are generated using incorrect degrees of freedom, which is often the case with thousands of non-independent SNPs (Do et al. 2014; Jones et al. 2016; Moran et al. 2019; Luikart et al. 2021; Waples et al. 2022). Finally, Waples (2023) also points out that adding more than a few thousand SNPs increases the precision only slightly and is more beneficial when the true N_e_ is large.

**Figure 3.**
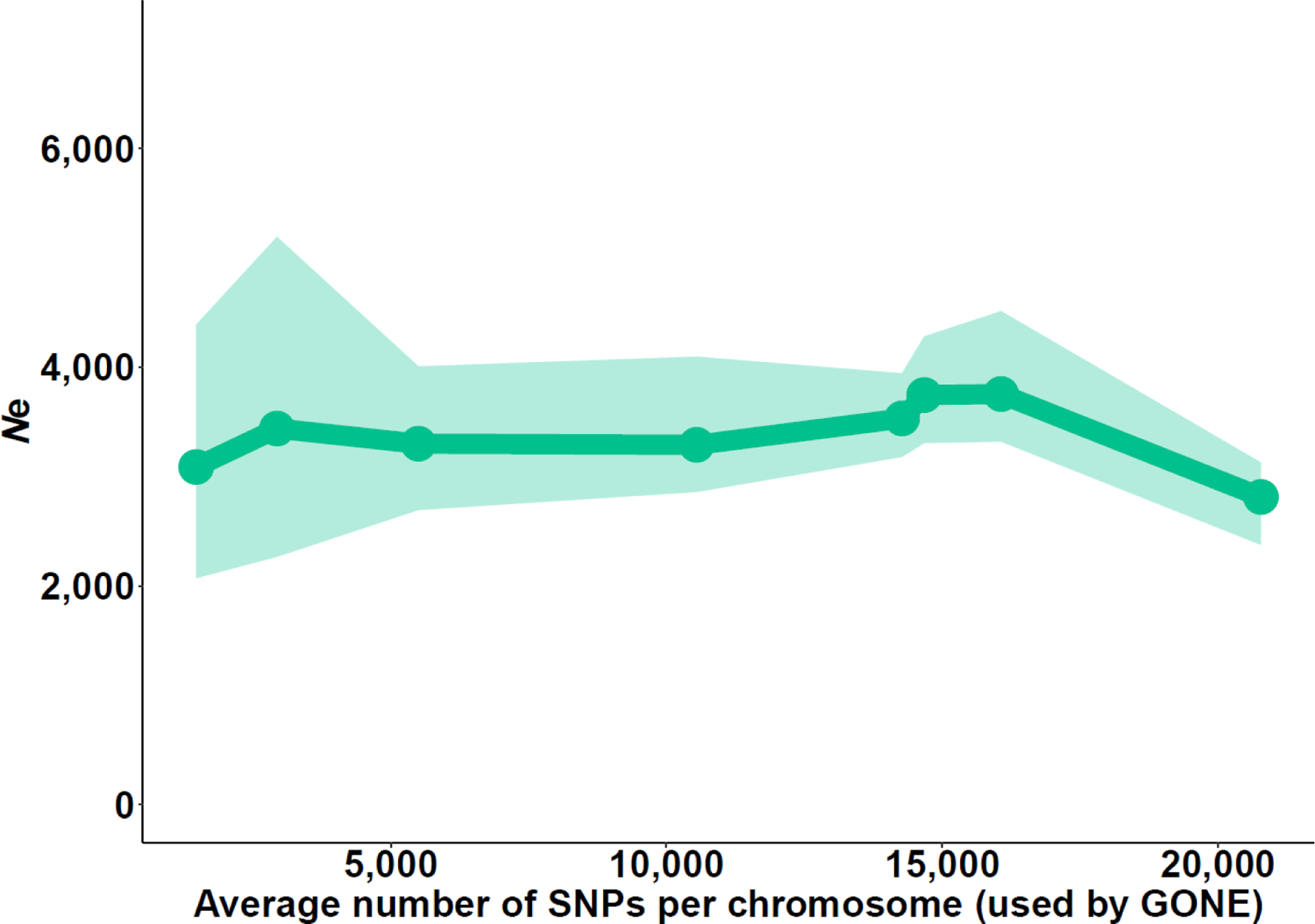
*N_e_* estimates obtained in GONE over the most recent generation for the Northern gene pool of *Prunus armeniaca* as a function of the number of SNPs. Points represent the geometric mean values across 50 replicates; shaded area represents 95% confidence intervals across replicates. Note that GONE uses a maximum of 50,000 SNPs per chromosome, even if provided with a larger number (with 1 million per chromosome being the maximum number accepted); the number of SNPs in each of the eight subsets analysed ranged from 10*^4^* to 10*^7^*, corresponding to a range of ∼5,000 to ∼20,000 polymorphic SNPs per chromosome used by GONE.

#### Influence of sample size on *N****_e_*** estimation

We evaluated the influence of sample size using the Northern gene pool of P. armeniaca. Increasing sample sizes to over thirty samples led to more consistent N_e_ estimates and reduced the chances of obtaining N_e_ estimates only representative of a few individual pedigrees (Fig. 4), as previously observed when using the linkage disequilibrium method (Palstra and Ruzzante 2008; Waples and Do 2010; Tallmon et al. 2010; Antao et al. 2011; Waples et al. 2016; Nunziata and Weisrock 2018; Marandel et al. 2019; Santiago et al. 2020). Including in the N_e_ estimation a number of samples that is representative of the true N_e_ of the population is crucial in large populations, where the genetic drift signal in recent generations is weak (Palstra and Ruzzante 2008; Luikart et al. 2010; Do et al. 2014; Barbato et al. 2015; Wang et al. 2016; Santiago et al. 2020; Waples 2023). On the contrary, small populations experience more genetic drift, hence the LD method is particularly powerful in such populations. Estimates of N_e_ remain small in small populations even with larger sample sizes, hence the important conservation implication that small populations cannot be mistaken for large populations (Waples and Do 2010; Waples et al. 2016; Santiago et al. 2020). For the Northern gene pool of wild apricots, we obtained an N_e_ estimate < 2,000 when sample size was equal to 15, and progressively obtained higher values increasing up to a plateau of N_e_ lll4,000, for larger sample sizes. This confirms the expectation that a large sample size is needed to estimate a large N_e_ (Tallmon et al. 2010; Antao et al. 2011).

**Figure 4.**
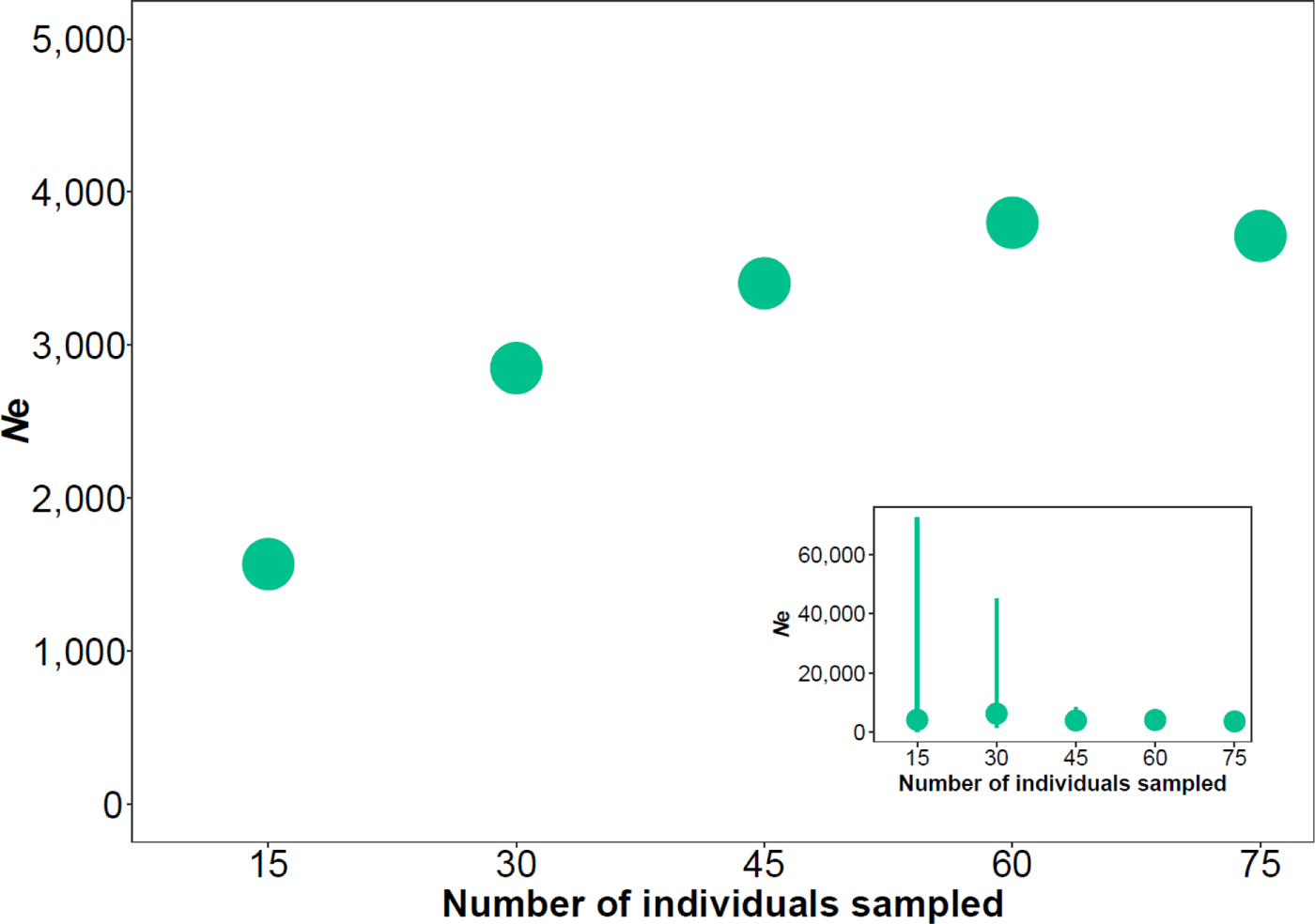
Change in the N_e_ estimates as a function of sample size in P. armeniaca (Northern gene pool). Points represent geometric means across subsets of individuals, sampled without replacement 50 times. The insert also shows 95% confidence intervals (point ranges) estimated over the 50 replicate subsets.

#### Influence of admixture on ***N***_e_ estimation

The impact of admixture on *N*_e_ estimation was explored using the dataset of *P. armeniaca*. Estimates of *N*_e_ in the most recent generation generally decreased when the Q-value of the individuals included in the analysis increased (Fig. 5A). The larger *N*_e_ estimates in the most recent generations (1-4) when including more admixed individuals are consistent with the upward bias predicted by Waples and England (Waples and England 2011) for a sampled subpopulation that does not include all potential parents (“drift LD”); with higher admixture proportions (Fig. 5A), the *N*_e_ estimated for each gene pool (subpopulation) using the LD method tends to approach the *N*_e_ of the metapopulation instead (Waples and England 2011). However, the *N*_e_ estimate we obtained when combining the two gene pools (“all” in Fig. 5A) was lower than the *N*_e_ estimate obtained when considering highly admixed individuals in the Northern gene pool (70% in the right panel of Fig. 5A). A downward bias in the *N*_e_ estimation is expected because of the Wahlund effect associated with sampling and analysing different gene pools together, and it is indicated as “mixture LD” (Waples and England 2011; Neel et al. 2013; Nunney 2016; Waples 2023). The Southern gene pool showed a contrasting trend; *N*_e_ estimates for the less admixed groups remained lower than that obtained when combining the two gene pools, possibly because the few samples from this gene pool contributed less (with any potential mixture LD) than the more abundant samples from the Northern gene pool (with their LD signal) (Fig. 5A). How the relationship between sampling and genetic structure practically affects *N*_e_ still deserves evaluation, as the effect on LDNe will depend on the relative strength of the “mixture LD” and the “drift LD” in the specific set of samples included in the analyses (Waples 2023).

Over the last 25 generations (Fig. 5B), we obtained higher *N*_e_ estimates when individuals from the Southern gene pool with a Q-value ≥ 99% were included. For the Northern gene pool, on the contrary, we obtained a lower *N*_e_ estimate when individuals with a Q-value ≥ 99% were included. The different demographic histories of the Northern and Southern gene pools certainly underlie the pattern observed, as the Southern gene pool seems to have undergone a recent bottleneck, whereas the Northern gene pool has a more stable demographic trend. The recent population decline for the Southern gene pool may be explained by the Soviet era and the current land-use change in the Fergana valley (mainly Uzbekistan) where native forests of wild apricot were partially replaced with crop species. Nevertheless, two more factors should be considered; first, the sample size of the Southern gene pool is smaller than that of the Northern gene pool (only 21 individuals vs. 77 individuals drawn from each Q-value subset). Second, Santiago et al. (2020) warn about a typical artefactual bottleneck observed in GONE and caused by population structure (in Figure 2F of Santiago et al. 2020, considering a migration rate = 0.2%; Novo et al. 2023). As we observed a consistent trend regardless of the individual Q-value, and the drop in *N*_e_ is particularly evident with a Q-value = 99%, we interpret this *N*_e_ drop as a true bottleneck, with the caveat of reduced accuracy linked to a small sample size for the Southern gene pool.

**Figure 5.**
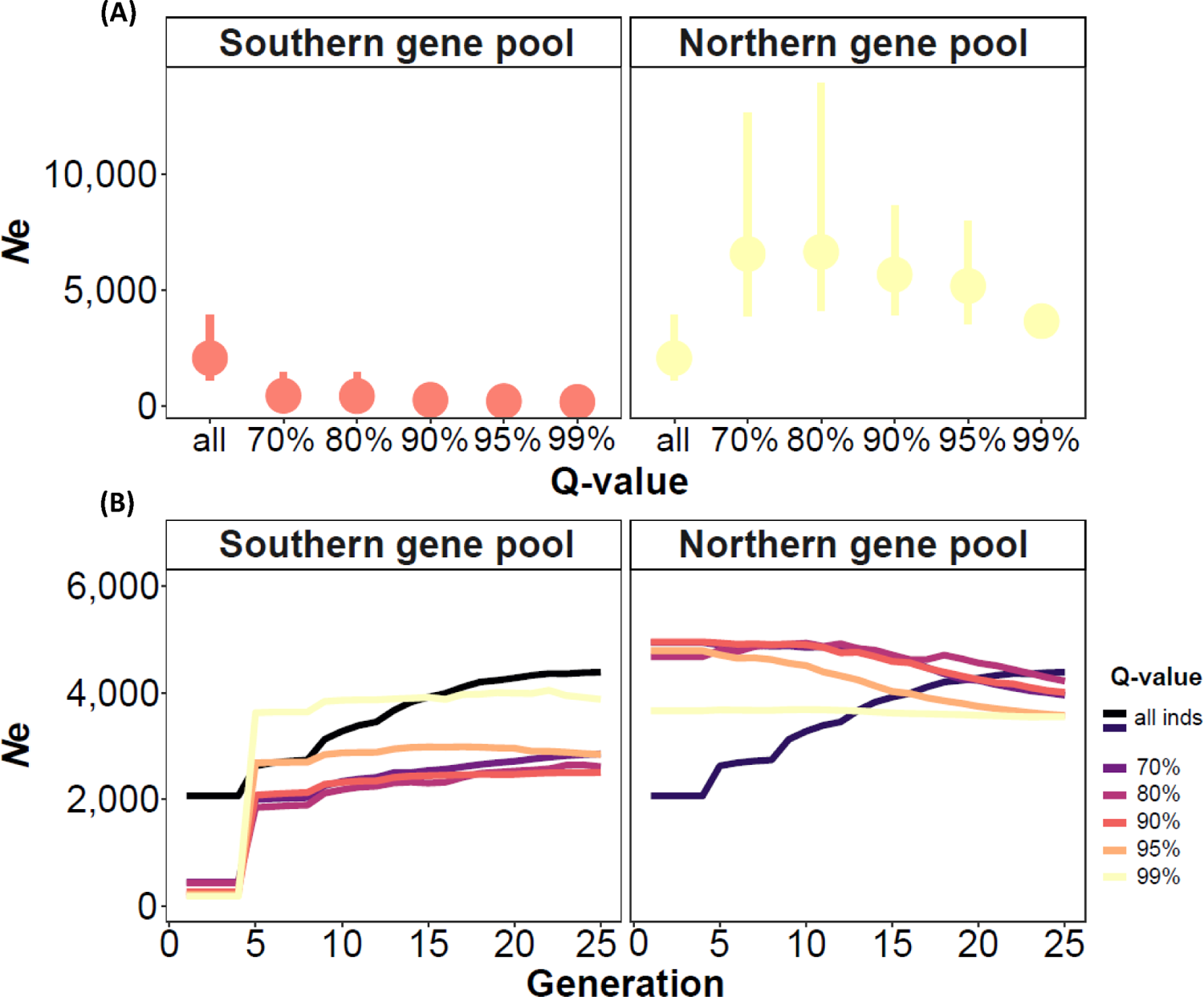
Influence of population structure on GONE *N*_e_ estimates for the Northern and Southern gene pools of *P. armeniaca* . Q-values refer to the results of the fastStructure analysis performed in Groppi et al. (2021) (lower bounds of individual Q-value to the main genetic cluster). *N*_e_ was estimated over 50 datasets of resampled individuals (77 in each Q-value subset in the Northern gene pool and 21 in each Q-value subset in the Southern gene pool, reflecting differences in sample sizes). In (A), points represent the geometric mean and ranges represent 95% confidence intervals across 50 replicates; in (B), only geometric mean values of the *N*_e_ estimates across 50 replicates and in the last 25 generations are shown. *N*_e_ estimates obtained for the combined gene pools are also shown (“all” in (A) and “all inds” in (B)).

#### Effect of using genomic scaffolds rather than chromosomes

To evaluate the effect of using genomic scaffolds as a proxy for linkage groups when chromosome information is not available, we sorted SNPs from the *P. armeniaca* dataset into a progressively larger number of scaffolds or chromosomes assumed. This produced inconsistent *N*_e_ estimates across the datasets with increasing number of chromosomes assumed, with *N*_e_ values progressively rising from around 3×10^3^ for 8 chromosomes (true value) to > 8×10^5^ when the number of chromosomes assumed was equal to 128 (Fig. 6). The algorithm implemented in GONE is based on the assumption that LD among pairs of SNPs at different genetic distances provides differential information about *N*_e_ at different times in the past (Santiago et al. 2020). Loosely linked loci give information about *N*_e_ in recent generations, as their recombination rate is higher and rate of LD-decay slower than that of closely linked loci (Sved and Feldman 1973). Therefore, the behaviour of the *N*_e_ estimates observed in Fig. 6 can be explained by considering that when a chromosome is broken into smaller scaffolds, only closely linked loci will be available for the *N*_e_ estimation; pairs of SNPs at higher genetic distances (i.e., loosely linked loci) will be missing, inducing biases on recent *N*_e_ estimates. An inflated *N*_e_ in recent generations will therefore depend on having fewer random associations among loci useful to estimate LD (i.e., fewer loosely linked loci), which will unfold as having less genetic drift (i.e., a larger population). Consequently, *N*_e_ estimates obtained in GONE for *M. annua* and *F. sylvatica* may be biased upward since scaffolds were used as a proxy for chromosomes (Table 1).

**Figure 6.**
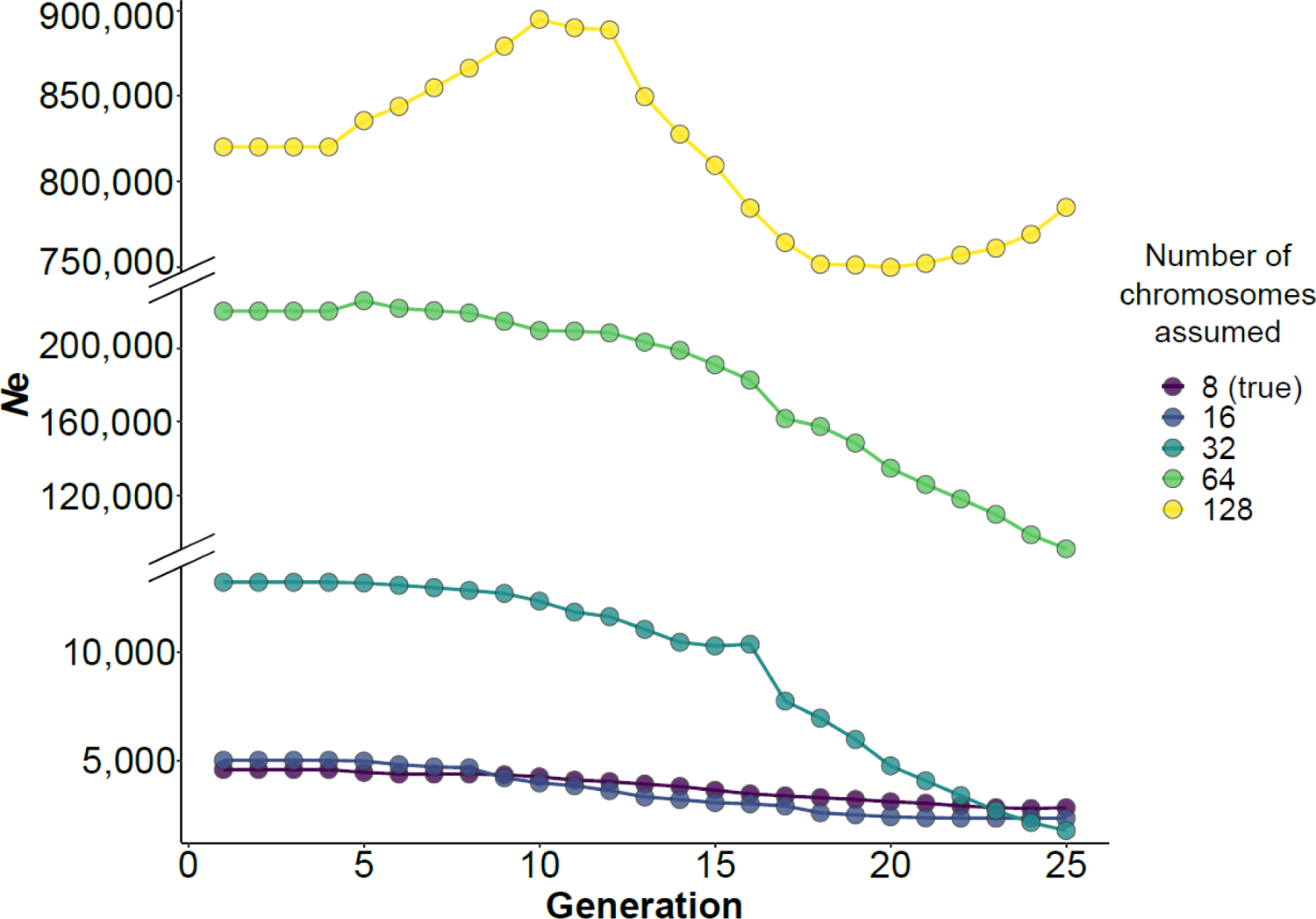
Estimates of *N*_e_ calculated on datasets in which the same set of SNPs is assigned to a progressively larger number of assumed chromosomes, where 8 is the true number of chromosomes for *P. armeniaca* (per haploid count); 45 individuals from the Northern gene pool were used for this analysis.

### N_e_ estimates obtained in GONE, NeEstimator and currentNe

As expected, *N*_e_ estimates obtained using NeEstimator and currentNe were more in agreement with one another compared with those obtained in GONE for the last generations (Table 2). GONE estimates for all species were larger than those obtained using the other programmes, especially in the Northern gene pool of *P. armeniaca* (GONE-*N*_e_ ∼3500 for the last generation while NeEstimator-*N*_e_ ∼716.2, excluding singletons and after bias correction, and currentNe*-N*_e_ ∼450 after bias correction). The point *N*_e_ estimate obtained in currentNe and its confidence intervals remained consistent even when we increased the number of SNPs included in the analysis, suggesting that there was no uncertainty associated with the SNPs included in the analysis. Estimates from simulated populations in Santiago et al. (2023) showed consistency between the output of currentNe and NeEstimator, except when a small sample (10 individuals) was drawn from a very large population (*N*_e_ = 10,000) using 22,000 SNPs, in which case currentNe performed better. Our sample size for the Northern gene pool was much larger (77 individuals), and we do not expect the true *N*_e_ to be larger than 10,000. Therefore, when using the same dataset for currentNe and NeEstimator, we interpret the slight discrepancy between the two estimates to be associated with the different algorithms included in the programmes, which are affected in different ways by the occurrence of rare alleles and the deviations from random mating, among other things (Santiago et al. 2023). When considering the Southern gene pool, for which the true *N*_e_ is expected to be smaller than for the Northern gene pool (Groppi et al. 2021), the estimates obtained in NeEstimator (∼80.9 excluding singletons and after bias correction) and *currentNe* (∼76.4 after bias correction) were more consistent.

**Table 2.**
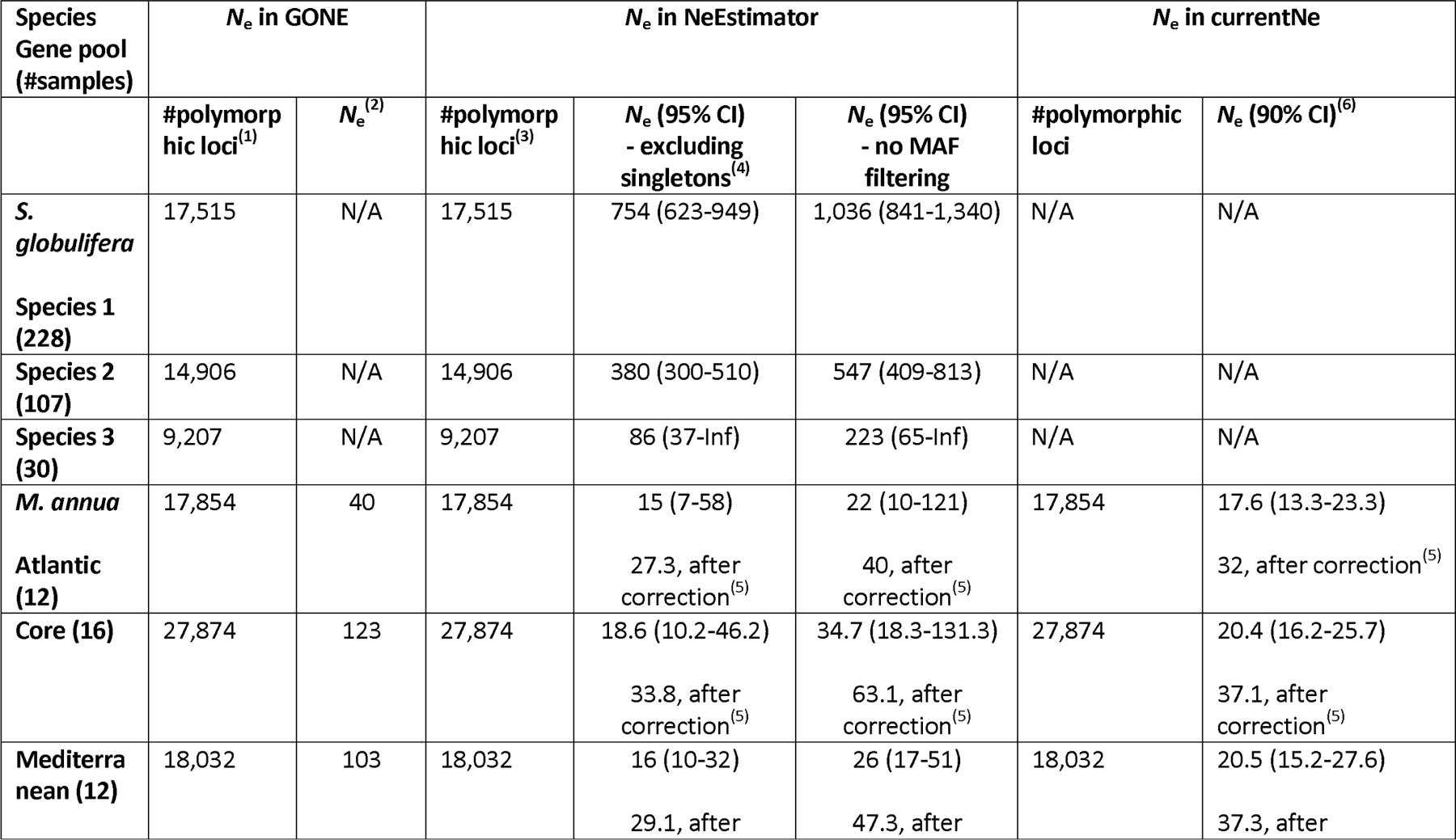
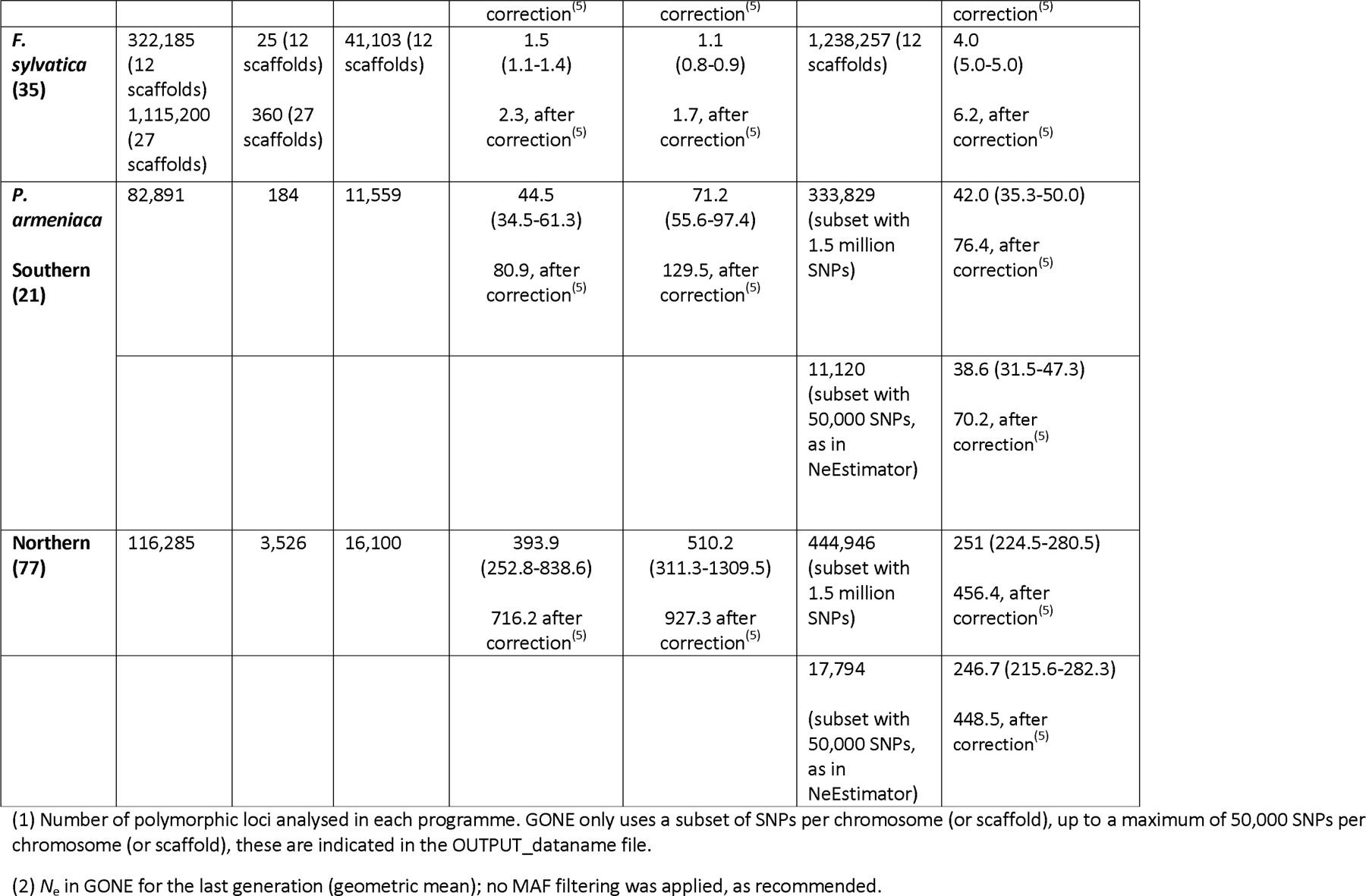
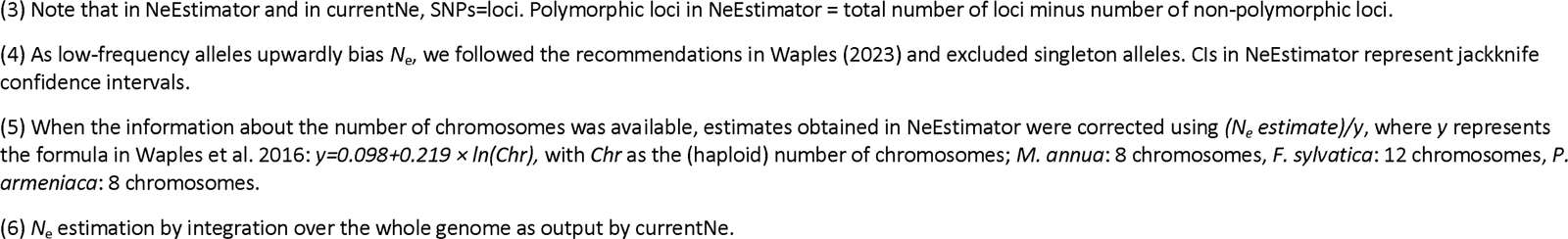
Estimates of effective population sizes for each dataset analysed in GONE, NeEstimator, and currentNe.

Another consideration is the downward bias on *N*_e_ estimates caused by localised sampling in continuous populations featuring isolation by distance (Neel et al. 2013; Nunney 2016; Santos-del-Blanco et al. 2022; Waples 2023). If the range of sampling is similar in extent to the unknown effective range of dispersal, as it is likely the case in *S. globulifera*, estimates may not reflect the population-wide true *N*_e_, but rather a quantity close to the neighbourhood size (*N*s), i.e., the inverse of the probability of identity by descent of two uniting gametes (Santos-del-Blanco et al. 2022). In *P. armeniaca*, where the sampling window likely exceeded the breeding window by much, we may still expect a downward bias because of the mixture LD caused by the inclusion of genetically divergent individuals (Neel et al. 2013; Waples and England 2011; Waples 2023). However, this bias would not explain the discrepancy between the estimates obtained in GONE and those obtained with the other programmes for the Northern gene pool of *P. armeniaca*. In *S. globulifera*, for which we also expect a large *N*_e_ (> 1000), it was only possible to use NeEstimator, due to the short length of contigs (not appropriate when using GONE), and the lack of information about the number of chromosomes (as required by currentNe). *N*_e_ ranged from 86 (CI: 37-Infinite) in Species 3, to 380 (CI: 300-510) in

Species 2 and to 754 (CI: 623-949) in Species 1, although point estimates could not be corrected for physical linkage due to lack of information about chromosome number and are therefore biased downward (Table 2). Estimates for Species 3, in particular, displayed infinite confidence intervals, suggesting that the sample size might be not large enough to capture the genetic drift signal from the original population. However, the relative magnitude of the estimates obtained are in agreement with the availability of suitable habitats for the three species (Schmitt et al. 2021) and, all else being equal, we would generally expect these populations to have a long-term constant population size, considering that the Guianese rainforest has experienced a continuous forest cover since the last glacial maximum (Barthe et al. 2016).

The uncertainty in *N*_e_ estimation using the LD method is particularly exacerbated in the dataset from *F. sylvatica*, where missing data also affect the estimation performed with the three programmes (GONE-*N*_e_ = 25 for the last generation, NeEstimator-*N*_e_ 12.3, excluding singletons and after bias correction for physical linkage, and currentNe-*N*_e_ 16.2 after bias correction for physical linkage), by reducing the usable sample size among pairs of loci (Peel et al. 2013; Do et al. 2014; Waples 2023). In general, missing data affect the precision of *N*_e_ estimates from the LD method whereas accuracy should be less affected (Nunziata and Weisrock 2018; Waples 2023), unless missing data occur non-randomly and depend on the genotype, as it might be the case in the *F. sylvatica* dataset.

For the only annual plant in our dataset, *M. annua*, we would expect *N*_e_ estimated with the LD method to mainly reflect the effective number of breeders, *N*_b_ (Luikart et al. 2021; Waples 2023) for the year of sampling, as individual cohorts were sampled (progeny of adults that reproduced in that specific year). Estimates in GONE were higher than those obtained in NeEstimator and currentNe (Table 2), also because of the bias induced by the lack of SNPs mapping (i.e., using scaffolds as a proxy for chromosomes in GONE). All point estimates fell within the estimated confidence intervals and usually denoted a small *N*_e_, which is consistent with primarily reflecting the *N*_b_ for the population. In particular, point estimates in NeEstimator, excluding singletons and after bias correction for physical linkage, ranged from 29.1 for the Mediterranean gene pool to 33.8 for the Core gene pool and 27.3 for the Atlantic gene pool. Point estimates in currentNe, after bias correction for physical linkage, ranged from 37.3 for the Mediterranean gene pool to 37.1 for the Core gene pool and 32 for the Atlantic gene pool. Even if the gene pool subdivision was consistent with the level of genetic admixture found in the individuals, it is still possible that estimates are biased downward because of mixture LD associated with mixing samples from different geographical locations (sampling window larger than breeding window). Furthermore, *M. annua* is able to survive through multi-annual seed banks (Crocker 1938) despite being an annual plant, and therefore the arithmetic mean across multigenerational *N*_b_ estimates would be needed to reliably estimate *N*_e_ rather than *N*_b_ (Nunney 2002; Waples 2006b).

### Practical recommendations when estimating contemporary N_e_ in GONE

In this study, we have considered some of the technical limitations when estimating *N*_e_ from plant genomic datasets, including: (i) the occurrence of missing data, (ii) the limited number of SNPs/individuals sampled, (iii) the lack of genetic/linkage maps and of information about how SNPs map to chromosomes when estimating *N*_e_ using the software GONE. In addition, we have explored some biological limitations that may affect *N*_e_ estimation using the LD method, such as the occurrence of population structure, although we recognise that our exploration is not exhaustive, as other biological factors (i.e., associated with reproductive system and life-history traits) might affect *N*_e_ and its estimation. Our empirical results corroborate some previous findings, for example about the importance of having large samples sizes (ideally > 30 per subpopulation), especially when populations are large, and highlight the following requirements that genomic datasets should satisfy:

⍰ non-random missing data should not exceed 20% per individual. Missing data also affect how SNPs are represented across loci and individuals sampled and can generate non-random patterns whose effect on *N*_e_ estimation is difficult to predict;
⍰ having a large number of SNPs (> tens of thousands) is potentially important to allow users to generate non-overlapping subsets of loci that reduce the influence of pseudoreplication on confidence intervals (Waples et al. 2022). However, increasing the number of SNPs beyond a few thousands per chromosome does not produce significant changes in *N*_e_ estimates, as we observed in wild apricots; Waples (2023) also observed that the benefit of adding over a few thousand SNPs on precision is little, but increases if the true *N*_e_ is very large.
⍰ most importantly, having SNPs fully mapped to chromosomes is essential to obtain reliable estimates when using the software GONE; other programmes should be preferred to estimate contemporary *N*_e_ when SNPs mapping is not available (i.e., currentNe).

In addition, the bias on *N*_e_ estimates due to the occurrence of gene flow and admixture can significantly affect the performance of single-sample estimators, as previously described (e.g., Neel et al. 2013). Other biases associated with (i) further sources population structure (i.e., overlapping generations, demographic fluctuations including bottlenecks, reproductive strategies causing variance in reproductive success, etc.) and (ii) further technical issues associated with sampling strategies and genomic datasets can add up and generate results that are misleading for conservation. Therefore, a careful consideration of the issues above is essential when designing and interpreting studies focused on the estimation of *N*_e_ and other related indicators for conservation.

## Data accessibility and Benefit-Sharing

The SNP matrices used in this study can be accessed at the following links: <u>10.5281/zenodo.4727831</u> (*Symphonia globulifera*), https://datadryad.org/stash/dataset/doi.10.5061/dryad.74631 (*Mercurialis annua*), 10.57745/FJRYI1 (*Fagus sylvatica*), <u>10.5281/zenodo.8124822</u> *(Prunus armeniaca*). The analyses carried out in this study and the related scripts are available at: https://github.com/Ralpina/Ne-plant-genomic-datasets (Gargiulo, 2023).

Benefits Generated: benefits from this research accrue from the sharing of our data and results on public databases as described above.

## Acknowledgements and Funding information

This study was carried out within the short-term scientific mission “Estimating effective population size in genomic datasets: test of methods and assumptions”, organised by Working Group 2 of the European Cost Action CA18134 “Genomic Biodiversity Knowledge for Resilient Ecosystems (G-BiKE)”. The work on *F. sylvatica* was supported by the Genoscope, the Commissariat à l’Énergie Atomique et aux Énergies Alternatives (CEA) and France Génomique (ANR-10-INBS-09–08). We are grateful to the Genotoul bioinformatics platform Toulouse Occitanie (Bioinfo Genotoul, <u>10.15454/1.5572369328961167E12</u>), to the Bordeaux Bioinformatics Center (CBiB), and to the Royal Botanic Gardens, Kew HPC (KewHPC) for providing computing and storage resources. We thank Enrique Santiago and Armando Caballero for their suggestions on how to interpret parameters and results using the software GONE and currentNe, and Stéphane Decroocq for the assistance with the wild apricot dataset. We thank Iris Biebach, Alice Brambilla, Christine Grossen, Jo Howard-McCombe, and all the other members of the G-BiKE Working Group 2 chaired by Mike Bruford for the useful discussions about *N*_e_ estimation methods and strategies. IP-V was supported by the U.S. Geological Survey Powell Center for Synthesis and Analysis.

