## Supplementary File 2 for "Estimation of contemporary effective population size in plant populations: limitations of genomic datasets"

#INPUT_PARAMETERS_FILE

########################################################

PHASE=2 ### Phase = 0 (pseudohaploids), 1 (known phase), 2 (unknown phase)
cMMb=1  ### CentiMorgans per Megabase (if distance is not available in map file).
DIST=1  ### none (0), Haldane correction (1) or Kosambi correction (2)
NGEN=2000  ### Number of generations for which linkage data is obtained in bins
NBIN=400  ### Number of bins (e.g. if 400, each bin includes NGEN/NBIN = 2000/400 = 5 generations)
MAF=0.0   ### Minor allele frequency (0-1) (recommended 0)
ZERO=1    ### 0: Remove SNPs with zeroes (1: allow for them)
maxNCHROM=-99  ### Maximum number of chromosomes to be analysed (-99 = all chromosomes; maximum number is 200)
maxNSNP=50000 ### Maximum approx number of SNPs per chromosomes to be analysed (maximum number is 50000)
hc=0.05   ### Maximum value of c analysed (recommended 0.05; maximum is 0.5)
REPS=40   ### Number of replicates to run GONE (recommended 40)
threads=30  ### Number of threads (if -99 it uses all possible processors)

###################################################################
